# The limitations of phenotype prediction in metabolism

**DOI:** 10.1101/2022.05.19.492732

**Authors:** Pablo Yubero, Alvar A. Lavin, Juan F. Poyatos

## Abstract

Phenotype prediction is at the core of many questions in biology. Prediction is frequently attained by determining statistical associations between genetic and phenotypic variation, ignoring the exact processes causing the phenotype. Here, we present a framework based on genome-scale metabolic reconstructions to reveal the mechanisms behind the associations. We compute a polygenic score (PGS) that identifies a set of enzymes as predictors of growth, the phenotype. This set arises from the synergy of the functional mode of metabolism in a particular environment and its evolutionary history, and is transportable to infer the phenotype across a range of environments. We also find that there exists an optimal genetic variation for predictability and demonstrate how the linear PGS can yet explain phenotypes generated by the underlying nonlinear biochemistry. Thus, the explicit model interprets the black-box statistical associations of the genotype-to-phenotype map and helps uncover what limits prediction in metabolism.

## INTRODUCTION

By understanding the factors that specify the phenotype, we aim to recognize how heritable biological information eventually maps onto action and how this mapping evolves (Waddington 2015). These issues illustrate the more general question of the emergence of function in complex systems, and the inherent attributes of these systems that help them elude function prediction (Orrell 2007). We will discuss here the properly biological case of metabolism.

The variational method of quantitative genetics characterizes a traditional approach to this question. Its goal is to establish statistical associations between the genetic and phenotypic variation observed within a certain population (Lynch and Walsh 1998). When this *genotype-to-phenotype* (GP) map becomes determined in a supervised situation, it is then possible to develop tools that infer the phenotype of individuals based solely on their genetic sequence (Dudbridge 2013). What are the limits of this approach?

Valid as they are, the statistical associations of quantitative genetics depend very much on the features of the trait, the population context, and the environmental conditions under which they are identified (Zaidi and Mathieson 2020). They thus represent a kind of “black-box” expectation that does not provide any insights into the processes causing a particular phenotype (Cannon and Mohlke 2018). This absence of mechanism has both basic and applied implications.

From the fundamental point of view, many features that define the genetic architecture of phenotypes (dominance, epistasis, etc.), while having a clear variational definition, present a less clear mechanistic interpretation (Keightley and Kacser 1987; Omholt et al. 2000). An interpretation that should also help explain how the nonlinearity that seems dominant in many biological systems limits the power of the –linear– statistical procedures (Feldman and Lewontin 1975).

From the applied point of view, consider, for instance, the case of genome-wide association studies (GWAS) in humans. The original purpose of GWAS was to identify the causal genetic determinants of complex phenotypes, including diseases. This plan turned out to be more complicated than expected (Visscher et al. 2017), with recent studies showing the complex pleiotropic regulation of most human traits (Boyle et al. 2017; Wray et al. 2018). Similarly, while specific prediction tools are indeed available, e.g., the development of polygenic risk scores to indicate a predisposition to disease (Torkamani et al. 2018), we still do not comprehend the biological foundations behind their successes and failures.

Indeed, discovering mechanistic insights behind these GP associations has proven to be a significant challenge, owing partly to the large quantity of accepted causal elements distinguished for most phenotypes. For instance, human quantitative traits were linked to only a few strong-effect determinants not long ago; a hypothesis that is now abandoned (Manolio et al. 2009; Boyle et al. 2017; Wray et al. 2018). A second factor is that natural selection weakens the impact of the *a priori* strongest statistical predictors (O’Connor et al. 2019). Most significant of all is the absence of an underlying developmental or physiological model explaining the emergence of the phenotype (Cannon and Mohlke 2018). Therefore, it is interesting to examine situations in which an explicit model replaces the black box so we can clarify the causality behind the associations.

There have been several attempts in this respect. Plant biology has pioneered works to connect gene network modeling with quantitative genetics, for example, on the prediction of flowering time (Welch et al. 2005). Other computational efforts to relate explicit phenotypic models and genetic variation include the cases of foliate-mediated one-carbon metabolism (Nijhout et al. 2017), single heart cells (Wang et al. 2012), or tooth development (Milocco and Salazar-Ciudad 2020).

In this manuscript, we employ a genome-scale metabolic model as a GP map to generate the genetic and phenotypic variation required to apply the methods of quantitative genetics. These models contain all the known metabolic reactions in an organism and the genes encoding each enzyme and enable the prediction of metabolic phenotypes, e.g., biomass, under situations in which genetics and environment can be controlled (Methods).

The approach has two advantageous features. First, we can induce genetic variation that does not suffer constraints experienced by natural populations, e.g., in the distribution of allele frequencies. We will thus be able to examine questions that depend on the details of the genetic variation. Second, and most importantly, it allows us to examine the processes driving the statistical associations and the potential of variational methods for phenotypic prediction.

Within this framework, we first discuss the concept of the polygenic score and then examine the underlying metabolic explanations behind its operation. For this, we will explore the influence of systems architecture, genetic variation, and gene-environment interactions. We show that the balance between the functional mode and the evolutionary history of metabolism is crucial in revealing the limitations of phenotype prediction with many implications.

## RESULTS

### Engineering genetic variation in metabolism

The genetic variation that exists in natural populations represents a multifactorial perturbation that enables us to understand biological processes (Rockman 2008). Quantitative genetics employs this perturbation to estimate statistical associations with phenotypes with the use of a reference, or training, population of known phenotype. The associations are quantified by the so-called polygenic scores (PGSs, Fig. 1A) that are eventually used to evaluate the effect of many specific genetic variants on an individual’s phenotype.

**Fig. 1.**
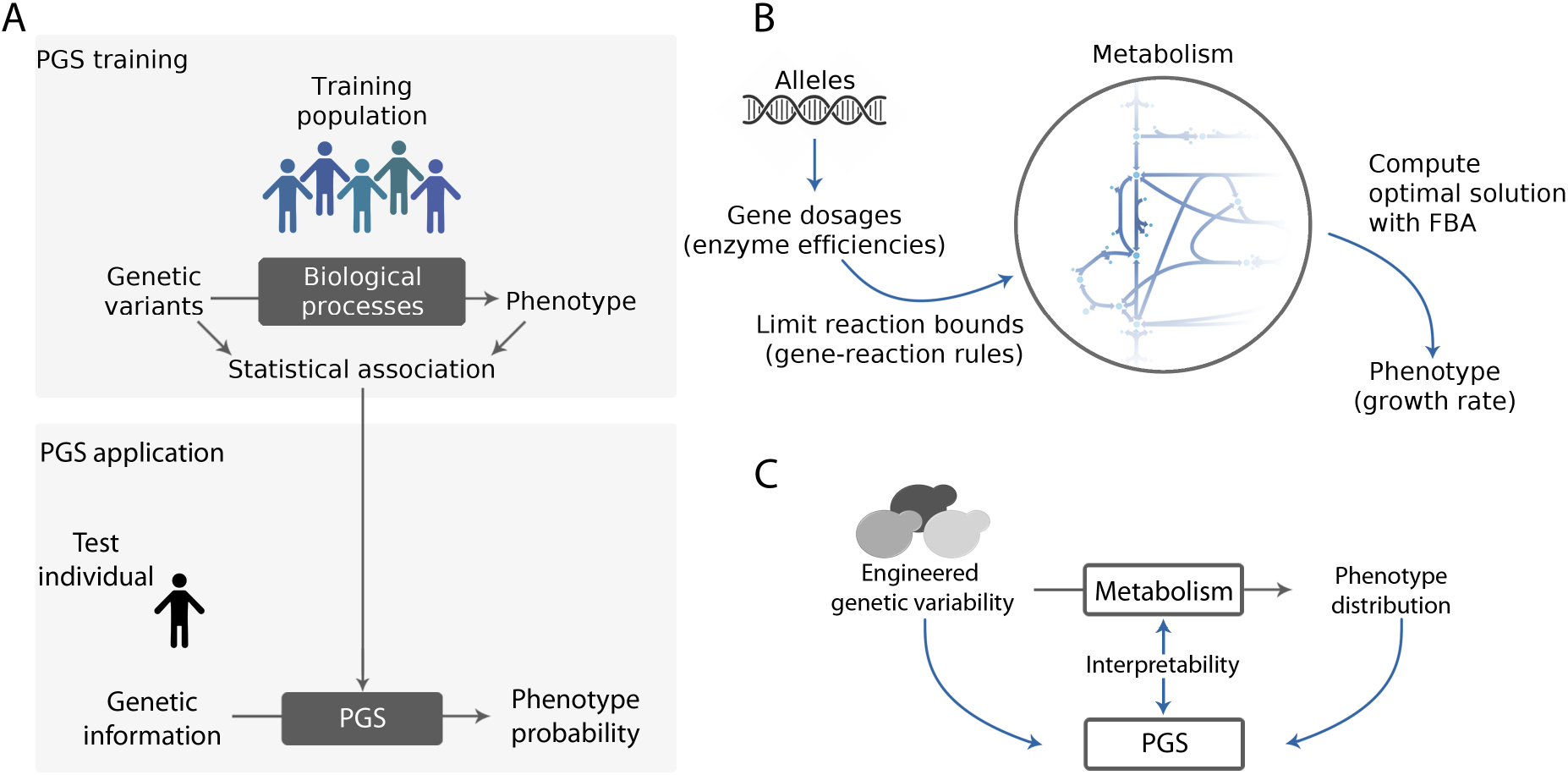
Metabolic reconstructions provide an explicit GP map that allows us to open the black box of phenotype prediction. (**A**) Phenotype prediction is typically based on the statistical association of genetic and phenotype variants in a training population. This approach defines a black box that bypasses all underlying biological processes. With the genetic information of a test individual, the model calculates a *polygenic score* (PGS) with the probability of observing the specific trait. (**B**) We benefit from the metabolic reconstruction of *S. cerevisiae* to generate an *in silico* population of yeast metabolisms. Genetic variation is modeled by the effect of alleles on gene dosages, which limits the maximum flux through their reactions according to the gene-reaction rules (Fig. S1 describes a specific example). Given these constraints, flux balance analysis (FBA) computes the growth rate of each individual in the population. (**C**) We quantify the statistical associations to elaborate a PGS for growth rate. Together with the availability of the underlying model, this enables us to “open” the black box of phenotype prediction to investigate its limitations.

Our first objective will be to *engineer* variation in the *in silico* metabolic framework. We consider whole-genome metabolic reconstructions and generate variation in gene dosages as a result of the genetic variation in the population. Dosages are relative to a maximal reference value resulting from the series of assumed different environmental and genetic contexts experienced by the yeasts, i.e., the evolutionary history of yeast metabolism (Fig. S1A). This reduced enzymatic performance is in line with earlier works (Kacser and Burns 1981; Keightley and Kacser 1987; Clark 1991) and are later interpreted quantitatively in the model by gene-reaction rules (GRRs). These rules –Boolean relationships between enzymes, Fig. S1BC-define which (and how) enzymes participate in reactions (Fig. 1B, Methods).

The genetic variation generated in this way originates individual differences in any potential metabolic trait, but we focus on the biomass production rate corresponding to the growth rate that is computed through flux balance analysis (FBA, Fig. 1B; see Palsson 2006 and the Supplementary Material for a concise introduction to FBA). Therefore, the entire procedure generates a data set of both genetic and phenotypic variations in the context of a metabolic model, which we can dissect to explain how the system works as a whole (Fig. 1C).

### A small subset of genes predicts growth within a metabolic polygenic score

We next derive a multidimensional PGS employing data of a population of yeast metabolisms that vary in their gene dosages as described before (growing in the standard medium, Methods). Different dosages induce variation in growth and in many metabolic fluxes (Figs. 2AB and Fig. S2). We obtain a PGS aimed at estimating the individual growth rate (Methods). Figure 2C compares the growth rate prediction to the values computed with FBA for this training data set. The PGS infers the phenotype with an 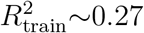, from here on abbreviated *R*^2^.

**Fig. 2.**
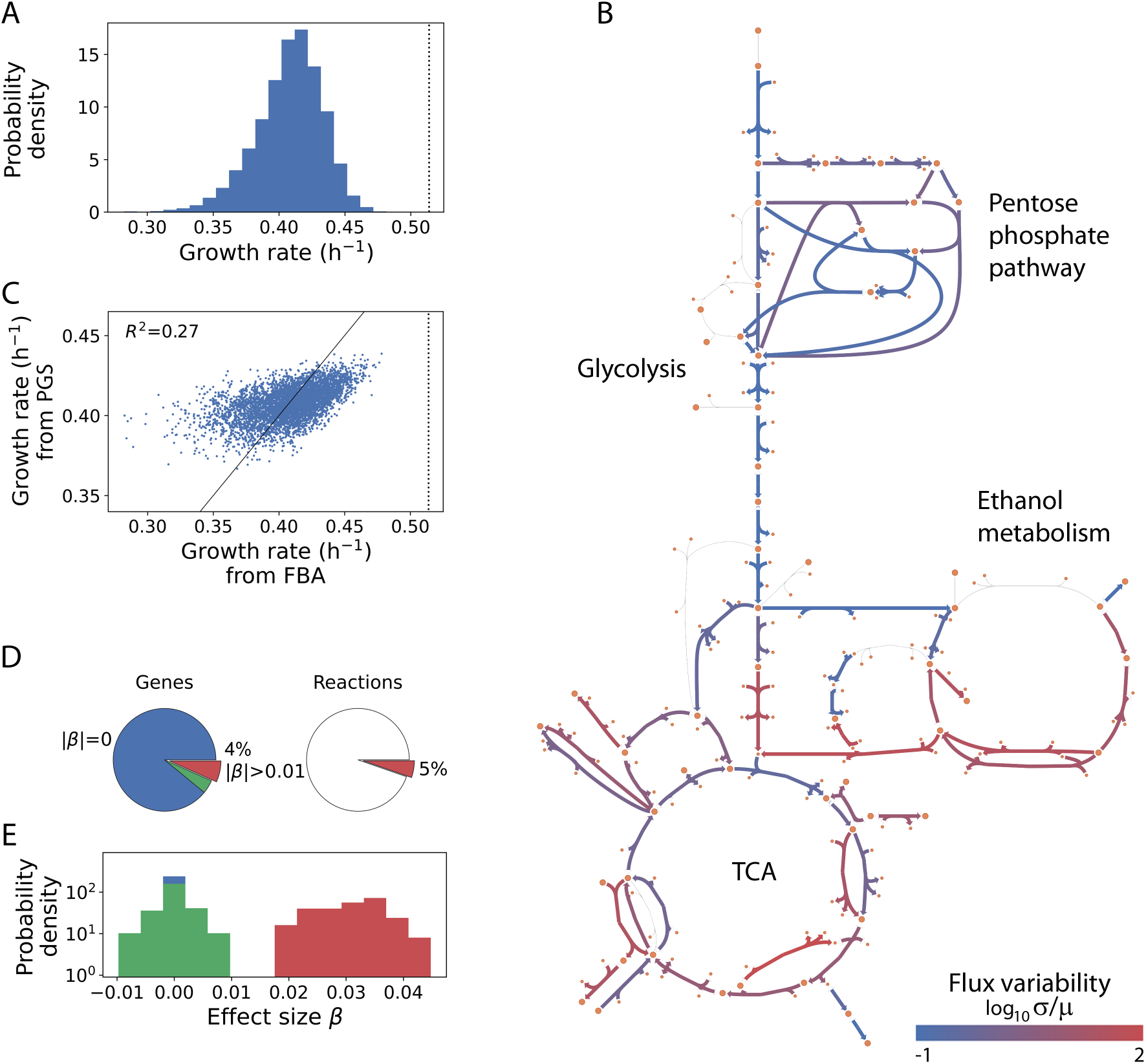
The PGS reveals that a small number of genes explains 27% of growth variation in yeast metabolism. (**A**) We generate a synthetic population of 5 × 10^3^ yeast metabolisms with relative gene dosages sampled from a normal distribution (Methods). For each individual, we compute its growth rate with flux balance analysis. Such genetic variation induces a distribution of growth rates with mean and deviation *μ* = 0.41 ± 0.03. The vertical dotted line shows the wild-type growth rate (*μ_wt_* = 0.51). (**B**) Central carbon metabolism scheme showing the variation in flux solutions across the population. (**C**) We trained a polygenic score (PGS) to predict growth rates from gene dosage data. The PGS explains 27% of the growth rate variation observed in the training population. (**D**) The PGS computes a specific effect size for each gene, *β*. Most of the genes, 88.7%, have null effect sizes (blue), while only 4.3% of the genes (red) are strongly associated with growth rate with |*β*| > 0.01. The latter control just 5% of the metabolic reactions. (**E**) Density distributions for each of the three categories before (null class is a delta distribution).

Although this situation differs from those typically observed in association studies –where the number of predictors is several orders of magnitude larger than the training population size–, it is exemplary in that we can easily overcome data shortcomings and hence the exact fit to the training data (statistical overfitting). That said, PGSs tend to lose predictive power when applied to a different *test* population. To evaluate this, we systematically generated 10^4^ independent test data, each with the same size and mutational distribution as the training population. For each test we computed the 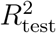 to obtain a mean 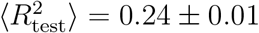 for the full set. This reveals that the computed PGS slightly overfitted the training data, and that we should expect a small loss of predictability when applied to different test populations 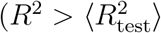 with *p* < 4×10^-3^.

The derived PGS includes all metabolic genes and their corresponding effect sizes, *β* (Methods). We identify 85 genes of non-null effect, 32 of which are comparatively large (|*β*| > 0.01; Fig. 2DE). The latter impacts 61 metabolic reactions (out of 1148). That most of the fluxes, in the considered standard medium, are inactivated explains this large number of genes with null effect (individual metabolisms within the population typically show ~73% of unused fluxes). Moreover, the set of active reactions characterizes a distinct metabolic “functional mode” (a relevant notion in what follows). From now on, we focus on the subset of genes with larger effect sizes.

### Few metabolic functions limit growth

What type of functions implement the predictor genes? One could think that predictors are distributed across all metabolic activities in the sense of universal pleiotropy (Kacser and Burns 1981). However, we only find a few metabolic subsystems enriched by predictors (Methods; Fig. S3). While these include the metabolisms of a variety of amino acids (valine, lysine, histidine), fatty acids, and phospholipids, it is surprising the absence of other subsystems central to metabolism like glycolysis or the citric acid cycle. These results are substantiated with a separate GO enrichment analysis (Table S1).

We observed that the detected subsystems involve specifically the production of biomass precursors. This group of metabolites fuels the biomass reaction, which defines the architecture of growth –as the trait of interest– in metabolic reconstructions (Fig. S4 shows its stoichiometry; this incorporates, for instance, the crucial role of amino acids and phospholipids for protein synthesis and the cell membrane, respectively, etc.). Consequently, we next hypothesized that the relevance of genetic predictors stems from their direct contribution to the pool of biomass precursors (Fig. 3A).

**Fig. 3.**
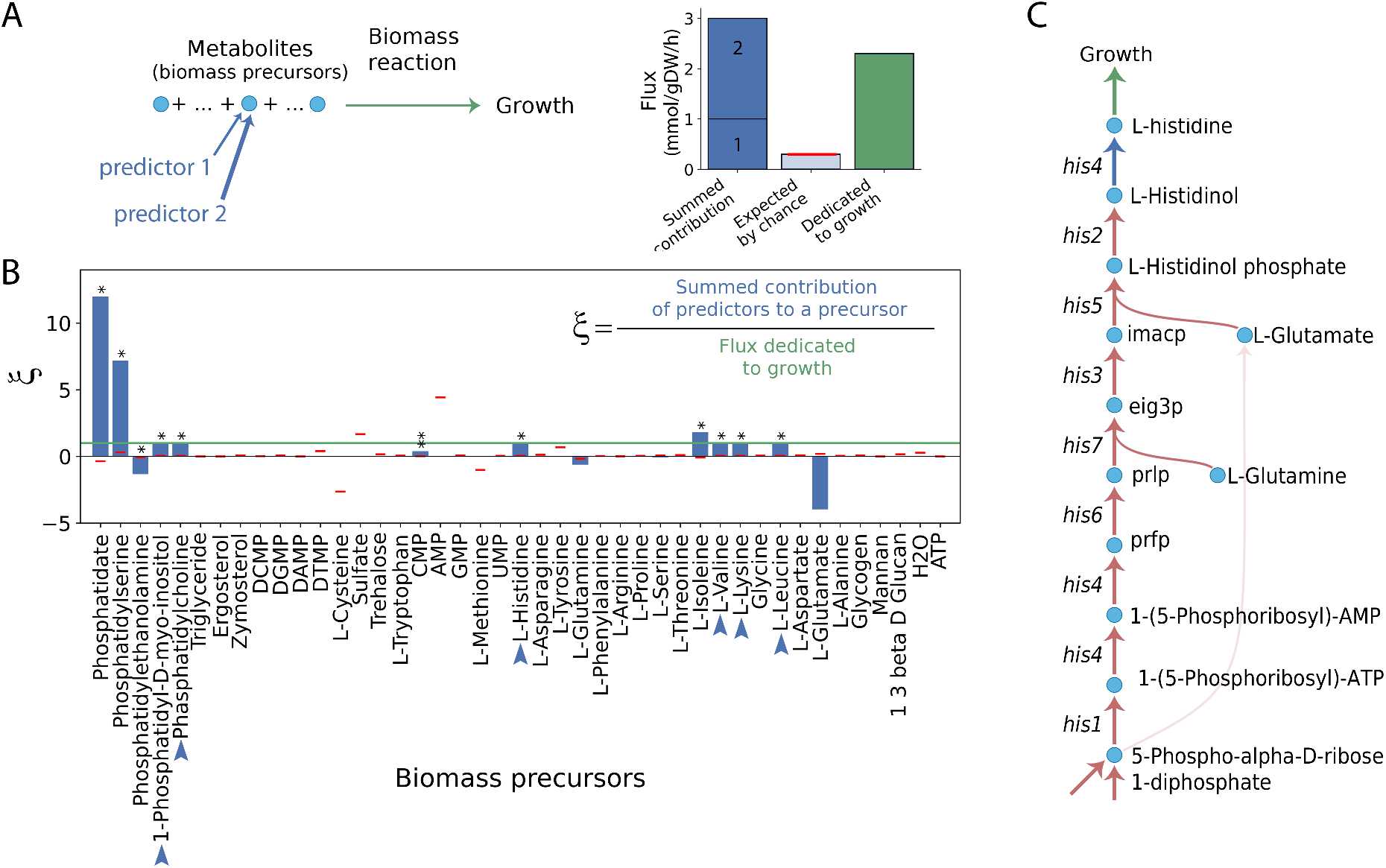
Top gene predictors control the production of a few metabolites required for growth. (**A**) Biomass precursors in metabolic reconstructions are metabolites that ultimately fuel the biomass reaction, which simulates growth. We compute the contribution of several genes (e.g., predictors 1 and 2, arrow line width describes each allocation) to the production of a specific precursor and compare it with the expected one of randomly selected genes (red horizontal line), or with the flux devoted to growth (green, this refers to the flux of the biomass reaction). (**B**) Mean values across the population of yeast metabolisms of *ξ*, the aggregate contribution of the predictor genes to the production of each precursor relative to biomass consumption. In some cases (blue arrows), the sum contribution of several predictor genes produces 100% of the precursor consumed by growth (horizontal green line). We tested significant contributions after 5×10^3^ gene randomizations controlling for subset size (mean, red horizontal lines; * p<0.05, ** p< 0.01). The case of L-glutamate is not significant due to a large variance (not shown for clarity). (**C**) Part of the metabolic pathway leading to L-histidine production. Although it is produced directly by *his4*, its production is influenced by upstream histidine-related genes, which also appear as important growth predictors.

Figure 3B shows the mean aggregate metabolite production (or consumption, if negative) associated with all predictors in the population of metabolisms (Methods; Fig. S2B and S5). The strongest predictors only contribute significantly to a subset of precursors (11 out of 43), and in some cases, e.g., valine, lysine, etc., this represents the total production that is required for growth. Therefore, the functional mode, active on the standard medium, effectively selects a domain within the entire architecture of growth. Of note, while the contribution to these factors is *direct* for only 9 genes –that are precisely producing these biomass precursors–, the rest of the predictors have an effect on growth in a somehow distributed manner.

The case of histidine is exemplary (Fig. 3C). While it is produced only by *his4*, all histidine-related genes are crucial to providing intermediary metabolites and thus are strongly involved in its overall production. This explains the case of *pmi40, sec53, dpm1* and *psa1*, which are predictor genes found upstream of the production of mannan, another important biomass precursor, while *pmt1-6* that ultimately produce said metabolite have null effect sizes. The same occurs with *erg4* and the production of sterol.

### Pleiotropy is not a good measure of growth predictor character

The results before underline that the top predictors include genes that directly alter the availability of the limiting biomass precursors and also genes whose impact comes through other upstream reactions. Could the systemic properties of metabolism capture this second aspect? We consider here first the pleiotropic character of a gene. One quantifies pleiotropy in metabolic models as the number of biomass precursors whose maximal production becomes reduced by changing the dosage of a gene (Shlomi et al. 2007). The score thus includes system-wide phenomena like metabolic compensation, rewiring, redundancy, etc.

Within the highly pleiotropic genes, only some display large effect sizes on the PGS (Fig. 4A). This outcome indicates that not all biomass precursors incorporated in the pleiotropic score limit growth in the standard medium, i.e., not all take part of the functional mode that is operating. Indeed, we confirm that our set of predictors are especially pleiotropic towards the group of biomass precursors already identified in the previous section, namely a few amino acids, phospholipids, mannan and sterol (Fig. 4B). This reflects that pleiotropy fails at pinpointing relevant genes for growth prediction since it is an aggregate measure that includes the effect of a mutation across *all* biomass precursors, while only a limiting few ones matter. On the contrary, separating the individual contributions of mutations to different functions results in a valid list of metabolites that potentially limit growth.

**Fig. 4.**
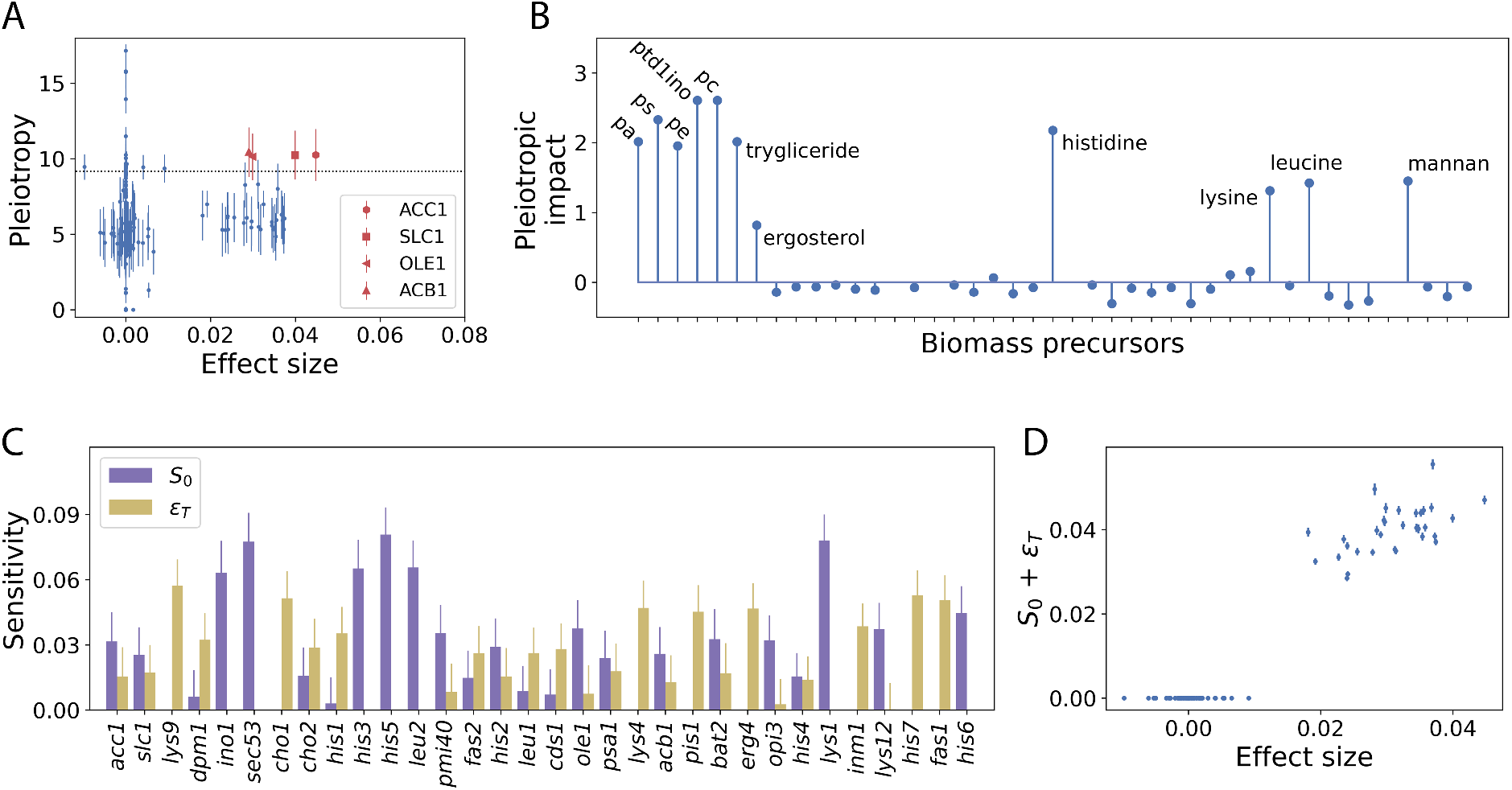
System-wide effects of top predictors. (**A**) Pleiotropy indicates the number of biomass precursors affected by a mutation of a given gene. Only a few top predictors have significant values of pleiotropy (over the 95th percentile of distribution of the pleiotropy for all genes, horizontal dotted line). Scores are computed considering a population of 10^3^ individual metabolisms (Methods). The plot shows the mean values (dots) and one standard deviation (error bars) across such population. (**B**) The pleiotropic impact of genes with large effect sizes focuses on a subset of biomass precursors. This is computed with the z-score of the sum of predictors (genes) that are pleiotropic w.r.t each metabolite against the mean and deviation found across all genes. We use abbreviations for phosphatidate (pa), phosphatidylcholine (pc), phosphatidylserine (ps), and phosphatidyl 1D myo inositol (ptd1ino). (**C**) Global sensitivity analysis allows us to quantify both the additive impact of genes on the growth rate, *S*_0_ (purple), and the total epistatic effects, *ϵ_T_* (yellow), which include 2nd and higher-order gene-gene interactions. Bars and vertical lines represent mean values and a standard deviation, respectively, of >10^6^ simulations (Methods). (**D**) The sum of the additive and epistatic effects correlates well with the effect sizes of the polygenic score (*ρ*>0.97, Pearson) demonstrating the validity of the global sensitivity analysis.

### Growth predictors display either large additive or epistatic effects

A second systemic measure is epistasis (indicating gene-gene interactions). Our objective is to understand to what extent the variation of the predictors contributes additively to the variation of the phenotype or if it also presents an epistatic component. To examine this, we introduce an approach based on *global* sensitivity analysis that precisely allows us to quantify the effect of individual gene doses on growth variation. Within this framework, two indices, *S*_0_ and *ϵ_T_*, estimate such additive and non-additive contributions (“total order” epistasis), respectively (Methods and Supplementary Material).

Figure 4C shows that some predictor genes have large additive effects, *S*_0_, and small total epistasis, *ϵ_T_* (*inol, his3*, …), whereas others display the opposite pattern (*cho1, his1*). Notably, the sum of all the effects, additive and epistatic, shows the maximum correlation with the effect sizes obtained in the PGS (Pearson’s *ρ*>0.97, Fig. 4D and Fig. S6) confirming the validity of the global sensitivity analysis. That the fraction of genes displaying *S*_0_>*ϵ_T_* and *S*_0_<*ϵ_T_* is comparable highlights that the large effect sizes we obtain are associated with genes enriched by additive effects (something to be expected from a linear statistical formalism) but also with those with strong epistatic effects (which appears paradoxical, see the Discussion; see also Figs. S7 and S8, and Supplementary Material for an alternative sensitivity analysis where the growth response coefficients are analyzed).

Is there a structural basis for large *S*_0_ or *ϵ_T_*? We investigate their relationship with several measures for each gene: the number of (active) reactions involved, the amount of flux they control, and the number of (precursor) metabolites they utilize. Among these, we find the largest correlations of *S*_0_ (and *ϵ_T_*) with the number of reactions they control, *ρ* = −0.19 (*ρ* = 0.19), and the log of the summed absolute flux through their reactions, *ρ* = 0.23 (*ρ* = −0.22). Therefore, one expects large *additive* effects to stem from genes that regulate a small number of reactions of larger flux. In opposition, genes that control a larger number of reactions with less flux result in larger *epistatic* effects.

### Optimal genetic variation for predictability

Once we have understood which metabolic and systemic aspects distinguish predictors in a fixed condition, we next investigate how changing these conditions influences predictability. Our first analysis focuses on examining the impact of the genetic variation available in the population, measured by its standard deviation σ_G_. We emphasize again that an advantage of our approach is that we can generate variation beyond what we might observe in a particular natural situation, where alleles frequencies could be restricted by the many evolutionary forces in play, e.g., natural selection, genetic drift, etc.

We use ten populations with equal mean dosage and increasing *σ_G_* (Fig. 5AB; Methods) to compute the corresponding growth rates with FBA and also to train a PGS in each case. We observe that the output of the PGS (coefficient of determination *R^2^*) reaches a maximal optimal value for a given *σ_G_* (Fig. 5B). This value results from stronger effect sizes for the same predictors found previously (Fig. 2D). Yet a better *R^2^* comes at the cost of a decreased mean population fitness (colored circles, Fig. 5B). In addition, the drop in PGS performance with higher genetic variation (*σ_G_* > 0.14) is due to the accumulation of new predictors with a strong effect resulting from the appearance of new growth-limiting reactions (Fig. 5C).

**Fig. 5.**
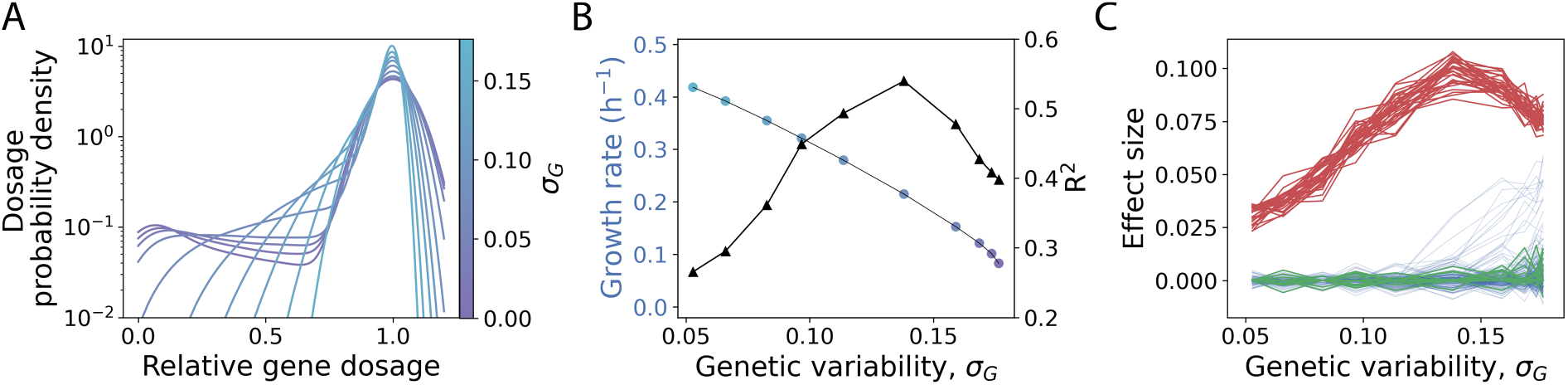
Populations with different genetic variation reveal a common PGS architecture but also its predictive limit. (**A**) Probability density function estimate (kernel, bw=0.3) of relative gene dosages of 10 populations with increasing genetic variation (*σ_G_*). (**B**) With FBA, we compute the growth rates (mean values on the left y-axis, colored circles as in panel A) and trained a separate PGS for every population (coefficient of determination *R*^2^ on the right y-axis, black triangles). (**C**) Effect sizes of all genes as a function of *σ_G_*. Lines are colored as in Fig. 2E.

That the effect of the common predictors changes as a function of the population genetic variation with which it has been estimated has important implications. When applying the PGS to individuals that show a specific set of alleles, in this case metabolic profile, the predicted phenotype will depend then on the estimation of the strength of the predictors, which, in turn, is dependent on the training set genetic variation. In sum, these results reveal a trade-off between genetic variation, population fitness, and predictability. While it is desirable to increase the performance of a PGS through sampling a population with high genetic variation, negative selection is likely to prevent scenarios of optimal predictability (O’Connor et al. 2019). Thus, selection would pose severe limitations to phenotype prediction.

### Prediction is portable across environments but also experiences extreme gene-environment interactions

In our final study, we asked to what extent specific growth conditions influence the ability to predict growth and at the same time modify the predictors. Therefore, we randomly generated >10^3^ (nutrient) environments of increasing richness (fixing the genetic variation as before, Methods). Then we train a separate PGS linked to the growth in every medium (Fig. 6A; we represent the different conditions as flasks with media of specific colors) to focus on the genes with the largest effect sizes (|*β*| > 0.01, as previously).

**Fig. 6.**
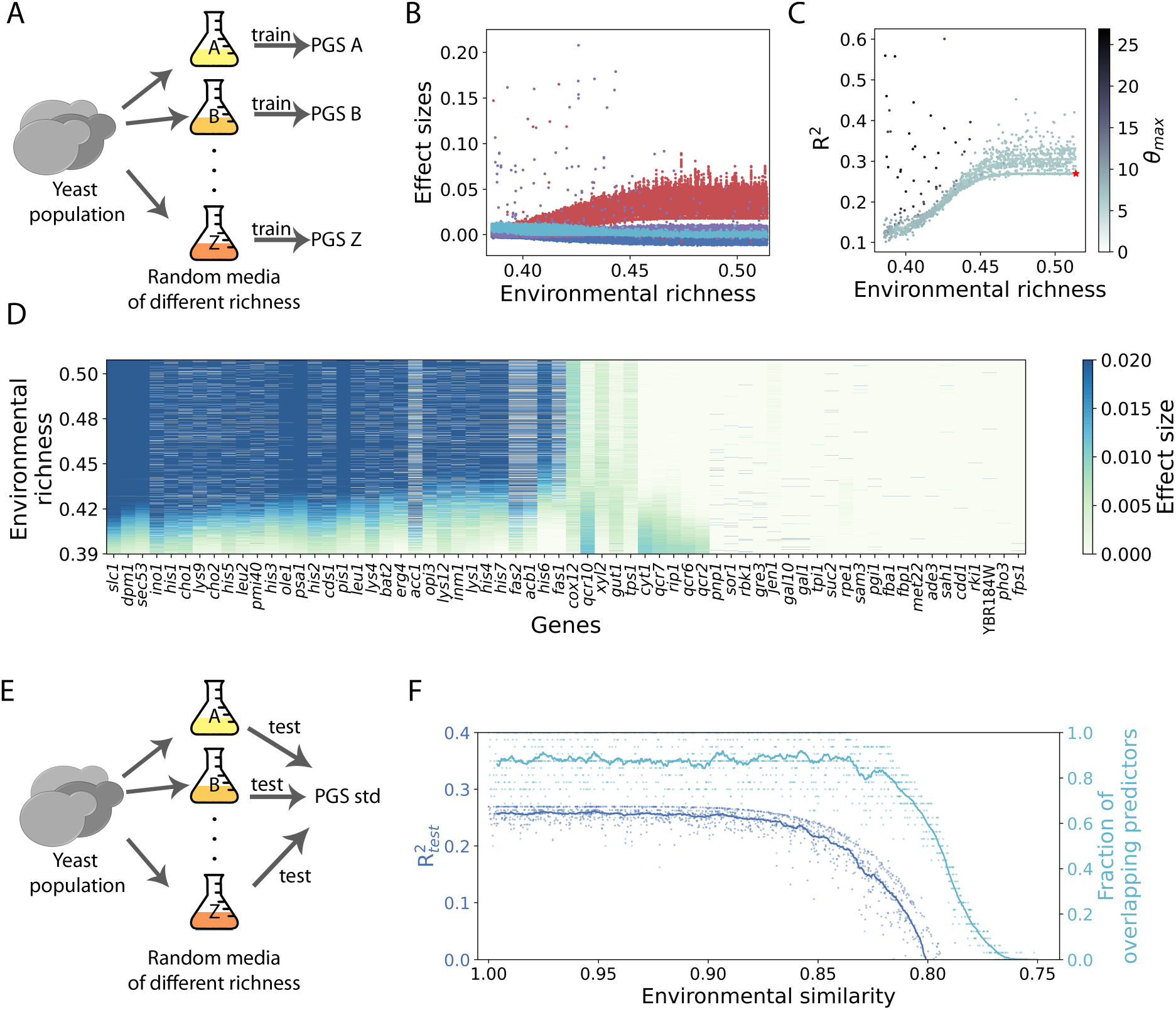
Environmental effects on phenotype predictability and the portability of a PGS. (**A**) We compute the growth rates of a fixed population in 10^3^ random environments (represented with flasks of distinct colors) with different richness (Methods) to train a PGS in each of these. (**B**) Effect sizes of all predictors forming the PGSs as a function of environmental richness. We highlight previously identified top predictors (red, as in Fig. 2), a novel set recurrent in poorer media that are related to the mitochondrial respiratory chain (cyan), and genes that show large effects in only specific media (purple). (**C**) The predictability of a PGS typically increases with environmental richness. However, in some media, predictability improves up to *R*^2^ = 0.6 due to strong gene-environment interactions identified by outliers in the effect sizes (*θ_max_* is the z-score of the strongest predictor found in each PGS, given its constituent effect sizes). (**D**) Effect sizes of genetic predictors follow a clear trend as a function of environmental richness. We show explicitly the values for all genes that have an effect size *β*>0.01 in any PGS. (**E**) Next, we test the “portability” of the PGS computed in the standard medium, PGS_std_, that is, its ability to predict growth rates in different environments. (**F**) The portability of PGS_std_ (left y-axis) holds within a certain environmental similarity, measured as the ratio of the random and the standard medium richness. The fall in “portability” is linked to the decreased overlap of predictors between PGS_std_ and the corresponding PGS of the medium (right y-axis). The dots and lines correspond to individual media and a running average (n = 50), respectively.

Figure 6B shows the effect sizes of each gene depending on medium richness. The top predictors identified earlier appear recurrently in most media, with increasing effect sizes in richer environments. Thus, they constitute a *core* set of predictors valid for most environments. Could this reflect that the growth medium has a limited impact on which reactions are active? That is, is the functional mode of metabolism similar? Indeed, there exist between 70% and 100% of shared active fluxes across all random environments, and >99% if we only consider reactions controlled by the initial set of top predictors. This explains the identification of similar genetic predictors across environments. However, we observe a second general trend in poor media, which exhibit a *different* functional mode, where a new set of predictors related to the mitochondrial respiratory chain becomes relevant.

One could still argue that the differences in predictability are due to subtle differences in metabolic solutions. We thus controlled for environmental richness to quantify this. Differences in metabolic solutions do not correlate with predictability (Pearson’s *ρ* = 0.06, using partial correlations to control for environmental richness), further pointing towards environmental richness as a valid measure that recapitulates metabolic activity and similarity, anticipating predictability (Fig. 6C).

Next, we observe that other genes recurrently appear as strong predictors in specific –typically poor– media (Fig. 6B), and whose occurrence leads to particularly strong PGS performance with up to *R*^2^ = 0.56 (Fig. 6C). Therefore, while growth prediction usually relies on a core set of genes largely “independent” of the growing medium, strong gene-environment interactions can sizeably improve the performance of a PGS.

Finally, the fact that the effect sizes change continuously with increasing environmental richness (Fig. 6B-D) ensures the “portability” of a singular PGS (trained in a reference medium) to predict the growth rate of the same population in another medium of similar richness (Fig. 6E). Figure 6F shows the performance of a PGS trained with data of the standard medium (PGS_std_) to predict the growth rates obtained in different random media (as test populations). Indeed, we observe that beyond a certain environmental similarity (Methods) the portability of PGS_std_ falls sharply together with the number of overlapping predictors between PGS_std_ and PGS_i_ (trained in the i-eth random environment). This enables us to distinguish regimes of high and low portability.

## DISCUSSION

We developed a framework to generate genetic and phenotypic variation in a population of *in silico* yeast metabolisms. The genetic variation considered does not in principle experience the restrictions that might be found in natural populations. Consequently, this method enables us to fully study how and why a PGS can infer the phenotype (growth rate) from a specific enzymatic profile given the explicit GP metabolic map that generates the phenotype (Kavvas et al. 2020). The quantitative interpretation of GRRs is a fundamental layer contributing to an accurate representation of this map.

In this way, if we reveal how the associations that define the PGS arise, we can begin recognizing general limitations of predictability, which have important implications for fundamental (Burns 1970; Boyle et al. 2017), and applied biology (Dudbridge 2013; Torkamani et al. 2018; Kavvas et al. 2020). Specifically, we evaluated the limitations originated by i) biological and evolutionary constraints of the metabolism, ii) knowledge of the trait architecture, iii) non-linearities in the GP map, iv) genetic makeup of the training population, and v) influence of the environment.

Computing a case study, we obtain an R^2^ = 0.27 and 32 genetic predictors with notably large effect sizes (Fig. 2DE). Which genes act as predictors result from the combination of two factors: the quantitative flux *required* –in a certain environment– in the reaction(s) associated with the given gene and the flux constraints *derived* from the corresponding genetic variation (Fig. S9A illustrates this combination). The former contributes to the “functional mode” of metabolism in that environment, while the latter results from the combination of GRRs and what we define as the “historical” reference flux bounds (see also Fig. S1). These bounds are a consequence of the adjustments of the yeast metabolism to both environmental and genetic conditions experienced during its evolutionary history.

Both GRRs and the reference bounds can act as a sieve of genetic variation causing part of it to be cryptic (Richardson et al. 2013; Paaby and Rockman 2014; Poyatos 2020). For instance, GRRs could describe the presence of an isozyme, and this redundancy would prevent such gene from becoming a phenotypic predictor. Genetic variation would also be cryptic when the reference bounds substantially differ from the required flux in a given condition. This again silences the functional impact of the genetic variation in the population. By manipulating reference fluxes, we confirmed this reasoning (Fig. S9B).

The presence of the 32 distinct predictors, and their enrichment in a few metabolic functions, is thus contingent on both working regime and evolutionary history. Note, however, that these functions should be necessarily associated with biomass precursors, either directly or in upstream reactions, since the biomass reaction represents the architecture of the trait/phenotype of interest, growth (Fig. 3AB).

Regarding what systemic properties characterize the predictors, we can report several results. Pleiotropy, as an *aggregate score* of the impact of mutations on all biomass precursors, is a poor measure of the predictive character of a gene, with the disaggregate information nevertheless partially revealing the composition of the biomass reaction (Fig. 4B). Therefore, the PGS provides a sound but partial understanding of the trait architecture, exclusively the domain associated with the necessary precursors in a particular environment.

We also discussed epistasis. Those genes whose dosages are eventually confining growth induce the non-linearities in the metabolic GP map, a combination of the functional mode and the evolutionary history as discussed above. On this basis, each individual is an instance where only one, or few, dosages are particularly limiting, the exact ones varying among them (Fig. S8). But the mixture of individuals at the population level generates functional cross-dependencies, increasing the number of limiting enzymes and consequently reducing predictability (more inadequate predictions correlate with predictor number, linear *ρ* = −0.94, Fig. S9C).

This interpretation helps clarify two added phenomena in which a reduced non-linearity enhances predictability. First, it describes the beneficial effect of sufficiently large gene-by-environment interactions on predictability (fewer predictors, better predictability, Fig. 6C). Second, it leads to a conjecture where a metabolism “disabled” on its capacity to react to genetic variation would paradoxically be coupled to better prediction. This premise, we proved (impairment causes simpler functional modes that lead too to fewer predictors, Fig. S9D). In short, the biochemical non-linearity of metabolic networks is related to the number of growth-limiting reactions active in the population, which limits the predictability of an individual’s phenotype.

Moreover, the validity of linear methods to predict the outcome of highly non-linear GP maps should be considered to be an enigma. Global sensitivity analysis confirms the suitability of these approaches (effect sizes capture the sum of both additive and epistatic contributions, Fig. 4D). Indeed, a PGS should capture non-linearities, since the minimization of error due to a linear regression incorporates all data points, including those coupled to non-linear regimes. Still, and despite the presence of epistatic effects, we notice that the sum of additive terms accounts for over 75% of the total phenotype variation ∑ *S*_0_ > 0.75, Fig. 4C). A signal that was also observed in a recent analysis of growth traits using a cross between two yeast strains (Forsberg et al. 2017). How is these additive component generated in our case? One of the previously discussed determinants is given by the extreme distribution of alleles in the population (Hill et al. 2008). A second determinant is conditional on the structure of this GP map, in particular on the monotonic relation between gene content and phenotype [order preservation of its responses to gene dosage, Fig. S8; (Gjuvsland et al. 2011)]. This monotonicity –particularly strong in the case of metabolic GP maps– is “broken” by the aforementioned functional cross-dependencies (see Supplementary material for further discussion and Fig. S9EF).

Finally, we asked about the portability of the predictions across populations concerning differences in genetic and environmental situations. Populations experiencing an intermediate genetic variation ensure maximum predictability, in line with previous results on extreme allele frequencies (Hill et al. 2008). However, such an increase in *R^2^* comes at the cost of population growth (Fig. 5B). Achieving this maximum could then be unattainable due to negative selection (O’Connor et al. 2019), which could be interpreted as yet another limitation on predictive power. In GP contexts in which genetic variation is less likely to cause loss of function and more forms of gain of function are possible, this constraint will be less apparent, as variation will not elicit negative selection.

Notably, the list of genetic predictors remains largely the same, independent of the amount of genetic variation in the training population (Fig. 5D). This is also observed when one determines predictors associated with populations growing in different media of similar richness, with two consequences (Fig. 6D). First, these results ensure the portability of the PGSs. Second, and as stated before, the predictors are specific to the evolutionary history of the metabolism. As a consequence, portability might not necessarily hold for more fixed, and maybe more realistic, histories.

In sum, we have learned that the combination of functional mode, evolutionary history, and phenotypic architecture determines the limits of prediction. For this, we benefited from an *in silico* approach whose capabilities to examine the emergence of phenotypic variation are beyond current experimental setups. Lastly, and while the principles learned here are general, in the sense that the structure and action of a metabolic GP map will show these properties, we still have much to study about maps associated with many other traits. The need to understand our predictions ensures many interesting future discussions.

## MATERIALS AND METHODS

### Metabolic models

Whole-genome metabolic models integrate the stoichiometry of the reactions in the metabolism of a model species, and together with computational methods they enable the estimation of an optimal network solution given an objective function where fluxes are stable. Among all fluxes, we focus on the prediction of biomass production, an analogue of growth rate and fitness. We used the genome-scale metabolic reconstruction of *Saccharomyces cerevisiae* iND750 (Duarte 2004) together with the Cobra toolbox for Python (Ebrahim et al. 2013) to compute the fitness of numerous mutants in either a standard medium or random media and the Escher package to depict the central carbon metabolism (King et al. 2015). Metabolic subsystems are typically assigned to reactions, hence we imputed a specific subsystem to a gene only if all reactions in which it participates belong to the same subsystem. Among all 750 genes present in the model, a subset of 42 had either none or multiple subsystems associated. We did not consider this subset when we compute Fig. S3. Note that our choice of model tries to balance the presence of sufficient biological details with accessible computational time. Still, our results and the mechanisms underlying phenotype prediction are robust when using the most recent yeast metabolic reconstructions iMM904 and yeast8 (*R*^2^ = 0.18 and *R*^2^ = 0.17, respectively).

### Quantitative mutations

We compute the effect of a quantitative reduction in gene dosage, or equivalently enzyme efficiency, in two steps (Fig. S1). First, we compute the wild type “reference” bounds of each reaction. These bounds are constituted by the maximum (and minimum if reversible) reaction fluxes observed in 2×10^4^ optimal solutions of metabolisms exposed to random environments (see next sections), and random genetic backgrounds. In the latter case, we randomly sampled flux bounds from a uniform distribution in the range [0,100] mmol/gDW/h. In this way, the wild type bounds integrate the history of yeast metabolisms, which have adapted to different environmental and genetic contexts (Discussion).

Second, we interpret quantitatively the gene reaction rules (GRRs) to find how reducing the dosage of an enzyme translates into a reduced flux through its reactions with respect to the wild type. This is necessary because some reactions may require several subunits or just one of several isozymes. The GRRs may contain AND and OR operators acting on pairs of genes, we consider these equivalent to min and sum, respectively, acting on relative gene dosages. This approach is similar to those used in noise propagation (Wang and Zhang 2011), or by the Escher package (King et al. 2015). In all cases, the upper/lower bounds are always computed and set according to the reactions’ reversibility and the bounds of ATP maintenance, biomass production and exchange reactions are kept unaltered.

This procedure is comparable to a previous approach in which genetic variation was also mapped to flux constraints (Kavvas et al. 2020). The authors construct the allele-to-flux constraint map coupled to the performance of a novel objective function to classify antibiotic resistance in a fixed medium. Our approach, however, assumes flux constraints imposed by a history of past genetic and environmental adaptations and, in this sense it is more comprehensive, together with explicit information about which genes are involved in exact reactions (GRRs). Finally, this method of associating genotypic variation with variation in fluxes, common in other previous works in the context of metabolism, e.g. (Keightley 1989), is a possible approach to the problem of how to characterize the genotype-parameter map for any model. The study of this map is a very stimulating research program in itself, but it goes beyond the focus of this study.

### Genetic variation

We generate genetic variation by sampling gene dosages from a probability distribution. Unless otherwise stated, we use a normal distribution with unit mean and standard deviation *σ* = 0.1. In this way, σ directly reflects the variation in the population’s gene dosages. This distribution follows from either Fisher’s original infinitesimal model, or from the Gaussian Descendants derivation where different levels of parenthood result in different *σ* under neutral evolution (Barton et al. 2017; Turelli 2017). We define the wild type genotype as having all gene dosages equal to the unit. This procedure generates populations that are in linkage equilibrium.

We also engineered genetic variation based on gamma distributions with shape parameters 0.5 < *k* < 200 and scale adjusted such that all distributions had equal mean. Importantly, note that gene dosages >1 are not beneficial, as wild type bounds are the extreme values observed (statistically), thus to avoid including additional cryptic genetic variation, gene dosages >1 were clipped to the unit. In this way, the resulting genetic variation in our standard population is *σ_G_* = 0.05 (Fig. 5).

### Growth media and environmental variability

The minimal medium is defined by unbounded import and export of H_2_O, CO_2_, ammonia, phosphate, sulphate, sodium and potassium and it is aerobic with 2 mmol/gDW/h import rate of O_2_. The standard medium is additionally composed of 20mmol/gDW/h import rate of glucose. Random environments are generated following a previous protocol (Wang and Zhang 2009). Briefly, we supplement the minimal medium with an additional number of components such that the probability of including any one follows an exponential distribution with mean *m* = 0.10 (other values produce similar results). Then, for every component, we obtain their maximum import rates from a uniform distribution between 0 and 20 mmol/gDW/h.

We define the richness of a medium as the growth rate of the wild type, and the environmental similarity between two media as the ratio of their richness. To avoid including arbitrarily rich media, we consider those with richness inferior, or equal to that of the standard medium. Also, we discard media that support biomass production rates <70% of that of the standard medium to avoid possible natural or model artifacts related to our implementation of quantitative mutations (see Results). This is an alternative approach to Constrained Allocation FBA (CAFBA), which limits the growth rate of metabolic models based on resource allocation principles by fixing a medium and tuning a parameter related to proteome fractioning (Mori et al. 2016).

### Polygenic Score

We used a high-dimensional regression framework for polygenic modeling and prediction:

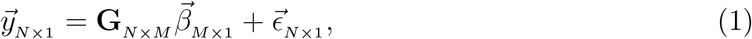

where *N* is the sample size, *M* is the number of genes, 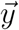 is the vector of phenotypes (typically growth rate), **G** is the genotype matrix, 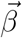 is the vector of effect sizes of the genes and 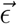 is some noise assumed normal with unknown variance. The generated data was fit using Least Absolute Shrinkage and Selection Operator (LASSO) a type of regression that under bayesian statistics assumes prior Laplace distributions in each coefficient, instead of uniform distributions as in the case of Ordinary Least Squares. Consequently, with LASSO some parameters are automatically zero (Tibshirani 1996), hence making it a remarkable alternative to pruning and thresholding (P+T) or other regularization methods (Dudbridge 2013; Wray et al. 2013). In addition, we compute the best value of the shrinkage parameter with five-fold cross validation. That effect sizes show a bimodal distribution makes our results robust to the application of other regularization, or feature selection methods (Fig. 2E).

### Sensitivity analysis and total epistasis

We computed local sensitivity indices *Z_i_* to monitor the changes in the output variable, i.e., growth rate, when every single input variable, i.e., each gene dosage, is altered (Kacser and Burns 1981; Keightley 1989). However, local sensitivity analysis (results are available in the Supplementary Material) does not explore the entire parameter space and is unable to isolate the effect of (non-linear) variable interactions.

Global sensitivity analysis, however, is ideal for this task, as it decomposes the variation of the output of a model into different terms when all variables fluctuate simultaneously. We used the method first proposed by Sobol for its easy implementation and interpretation (Sobol 1993; Saltelli et al. 2008). Note that this differs from previous flux-based applications (Nguyen Quang et al. 2019; Nobile et al. 2021). Briefly, we focus on two indices for the i-eth gene, the first order index 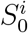 and the total effect index 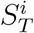. The former quantifies the additive part of the variation associated to a gene while the latter quantifies its total contribution, additive and all non-additive effects. From these, we derive the total epistasis which accounts for all, and only, non-additive effects as 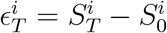 and its error 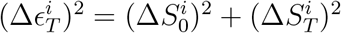.

We computed all indices and their errors with monte carlo estimators using over 10^6^ samples (Saltelli et al. 2008; Saltelli et al. 2010). We carried out these computations with pairs of genotypes sampled from the original population growing in standard medium. A detailed description of the protocol and equations are available in the Supplementary Material.

Note that we do not show negative values of both *S*_0_ and *ϵ_T_* as they are unrealistic and should be considered null in agreement with their error bounds.

### Pleiotropy

In metabolic models, the pleiotropy of a mutation is generally computed as the number of biomass precursors whose maximum production is limited by the mutation, following a previous protocol (He and Zhang 2006; Shlomi et al. 2007). In short, we simulated the excretion of a given metabolite by adding an exchange reaction to the model and maximizing the flux through this reaction. Then we consider that a gene limits the production of a metabolite if, when knocked-down by 90%, its excretion rate decreases. As pleiotropy is strongly dependent on the genetic context, we computed the mean value across 10^3^ individuals of the population due to the large computational load. We used a 90% decrease in dosage to avoid artifacts derived from gene essentiality, but our results are robust when using other values.

## DATA AVAILABILITY

Data and code for this work is available at Zenodo (Yubero 2022). The main code used to generate quantitative mutations is available at GitHub (https://github.com/pyubero/quantitative_mutations).

## ACKNOWLEDGEMENTS

This work was supported by grant PID2019-106116RB-I00 (P.Y. and J.F.P.), partially by Ph.D. fellowship BES-2016-079127 (P.Y.), and the program Severo Ochoa Center of Excellence (A.A.L.) from the Spanish Ministerio de Ciencia e Innovación and the European Social Fund.

## AUTHOR CONTRIBUTIONS

P.Y., A.A.L., and J.F.P. conceived and designed the study. P.Y. conducted all analysis with contributions from A.A.L. and J.F.P. P.Y and J.F.P. discussed the results and wrote the manuscript. All authors reviewed and approved the final manuscript.

## COMPETING INTERESTS

The authors declare no competing interests

## MATERIALS & CORRESPONDENCE

Correspondence and request for materials should be addressed to J.F.P.

## Notes

https://github.com/pyubero/quantitative_mutations

https://zenodo.org/record/6550708

### Competing Interest Statement

The authors have declared no competing interest.

### Summary of Updates

This version includes an extended discussion on the relationship between additive genetic variation, the monotonicity of metabolic genotype-to-phenotype maps, and predictability.

