## Supplementary material for "The limitations of phenotype prediction in metabolism"

### Supplementary Material
#### The limitations of phenotype prediction in metabolism

Pablo Yubero, Alvar A. Lavin, Juan F. Poyatos.
Logic of Genomic Systems Lab (CNB-CSIC) Madrid.

#### S1 Flux Balance Analysis

Flux balance analysis (FBA) is a computational technique that allows the simulation of metabolism from information about its structure. This information is now available for many organisms through genome-scale reconstructions of metabolic networks. Networks are represented in this formalism by a stoichiometric matrix of metabolites and reactions. Using computational methods, FBA calculates an optimal solution which comprises the quantitative values of the fluxes through all reactions. Such a solution is obtained by optimizing an objective function, which in most cases is the maximization of biomass production. Prior to this calculation, the flows of these reactions can be restricted, which ultimately delimit the space of possible solutions influencing the predicted phenotype. Also, nutrient exchange reactions are included in metabolic models and their bounds define the growth medium of the metabolism. See (Orth et al., 2010) for a primer description of this formalism.

#### S2 Local sensitivity analysis

Previous works on metabolism resorted to local sensitivity analysis to assess the impact of a model variable on any given metabolic flux (Kacser and Burns, 1981; Kacser et al., 1995). Analogously, we here compute the "control parameters" $Z_i$ of the i-eth gene on growth as

$$Z_i = \frac{\Delta \ln(\mu)}{\Delta \ln(g_i)}, \tag{1}$$

where $\mu$ is the growth rate in $h^{-1}$ and $g_i$ is the dosage of the i-eth gene relative to the wild type (Methods). Sensitivity values $Z_i$ quantify the individual (normalized) impact that each gene has on biomass production. As this measure is strongly dependent on the genetic context, we compute its average value across the entire population. Figure S7 shows that the control parameters of most genes (98%) are null, and only for a few $Z_i > 0$. These genes with non-null $Z_i$ are precisely those with large effects in the polygenic score (PGS).

Moreover, for a better understanding of the implications of large $Z_i$, we directly simulated the growth rate of >$10^2$ individuals when varying individual gene dosages from $g_i = 0$ to $g_i = 1$, while keeping their genetic background constant. The large response coefficients of the best predictors reflect that they are all strong limiting factors of growth under the range of dosages found in the population (recall that they follow a normal distribution with unit mean and $\sigma = 0.1$). Whereas the rest, with low $Z_i$, become limiting in only very rare events (Fig. S8).

### S3 Global sensitivity analysis

Global sensitivity analysis (GSA) is a mathematical toolset that contributes to the interpretability of complex models by decomposing an output variance into partial variances of input variables, or subsets of these. These methods have rapidly grown and have been widely used, for example in risk and model assessment. They have also been previously used in the context of flux balance analysis, but to assess the contribution of reactions instead of genes (Nobile et al., 2021). We summarize in the following the main ideas behind Sobol's approach to GSA and the protocol that we used (Sobol, 1993, 2007; Saltelli et al., 2008, 2010).

Imagine any model that takes $p$ input parameters $\vec{x} = \{x_i\}_{i=1}^p$ and outputs a scalar value $y = f(\vec{x})$. Here parameters and variables can be considered equivalent. Then the variance of the output $\mathrm{Var}(y)$ can be decomposed as:

$$\mathrm{Var}(y) = \sum_{i=1}^{p} V_i + \sum_{i=1}^{p} \sum_{j>i}^{p} V_{ij} + \ldots + V_{\vec{x}}, \tag{2}$$

which, dividing all terms by $\mathrm{Var}(y)$, can be simplified and rewritten as a function of the Sobol *indices S*:

$$1 = \sum_{i=1}^{p} S_0^i + \sum_{i=1}^{p} \sum_{j>i}^{p} S_{ij} + \ldots + S_{\vec{x}}. \tag{3}$$

This decomposition is particularly revealing as $S_0$ are the first order, fractional, contributions of each individual parameter to the total variance, while the rest include contributions of gradually increasing order, i.e. interactions between pairs of parameters, then triplets, etc. Apart from $S_0$, the total effects index is of particular interest, and it can be written as:

$$S_T^i = \sum_{\vec{w}_i} S_{\vec{w}_i}, \tag{4}$$

where all $\vec{w}_i$ contain the i-eth parameter. In this way, $S_T^i$ quantifies the fraction of variation associated to the i-eth parameter and all of its interactions with other parameters.

To compute these values, common approaches include Monte Carlo estimates and Fourier amplitude estimate testing (Saltelli et al., 2008). We focus on the former due to its simplicity and satisfactory convergence. Among different Monte Carlo estimators (Saltelli et al., 2010), we used the following for $S_0$ and $S_T$:

$$S_0^i = \frac{1}{N\,\mathrm{Var}\,(f(A))} \sum_{k=1}^{N} f(B^k)\left(f(A_{Bi}^k) - f(A^k)\right), \text{ and} \tag{5}$$

$$S_T^i = \frac{1}{2N\,\mathrm{Var}\,(f(A))} \sum_{k=1}^{N} \left(f(A^k) - f(A_{Bi}^k)\right)^2, \text{ with} \tag{6}$$

$$\mathrm{Var}\,(f(A)) = \frac{1}{N}\sum_{k=1}^{N} f(A^k)^2 - \left(\frac{1}{N}\sum_{k=1}^{N} f(A^k)\right)^2 \tag{7}$$

where $k$ is the sample, $N$ is the total number of samples, $f(A^k)$ is the growth rate of genotype $A^k$, $f(B^k)$ is that of genotype $B^k$, and $f(A_{Bi}^k)$ is that of genotype $A^k$ but with the dosage of the i-eth gene taken from genotype $B^k$. Also, A and B are genotypes sampled from our default population.

Therefore, the Monte Carlo protocol can be summarized in the following steps per sample:

1. Obtain two genotypes $A = \{g_i^A\}_{i=1}^l$ and $B = \{g_i^B\}_{i=1}^l$, where $l$ is the number of genes.
2. Create $l$ new genotypes $\{A_B^i\}_{i=1}^l$ such that all dosages are from A except for the i-eth which is taken from B, that is $A_B^i = (g_0^A, g_1^A, ...g_{i-1}^A + g_i^B + g_{i+1}^A + ... + g_l^A)$.
3. Compute $f(A)$, $f(B)$ and $f(A_B^i)$ for i=0,...,l.
4. Use Eq.(5) and Eq.(6) to compute $S_0$ and $S_T$, respectively.

### S4 Additivity on genotype to phenotype metabolic maps

Our goal here is to clarify the apparent contradiction between the substantial fraction of additivity found in the metabolic genotype-phenotype (GP) map and the relatively small values of $R^2$ of PGSs. According to Gjuvsland et al. (Gjuvsland et al., 2011, 2013), the monotonicity, or order preservation, of a GP map leads to a significant fraction of additive variation that should favor predictability (large $R^2$). We argue that the $R^2$ values we found are the result of a trade-off between the order-preserving nature of the GP metabolic maps (Gjuvsland et al., 2011) and the substructure of the population in terms of which genes are “predictors” in each individual (Hill et al., 2008).

To start, we observe that the monotonicity of the metabolic reconstruction is highly additive, i.e., order-preservation is not "broken" in any case [dosage-response profiles in Fig. S8 show a degree of monotonicity $m$=1, following (Gjuvsland et al., 2013)]. In addition, using global sensitivity analysis, we demonstrate that the sum of the first-order indices represents ~75% of total variance. Therefore, our model has a large fraction of additive variance. Why, then, is $R^2 = 0.27$ "only"?

First, despite displaying full monotonicity, our GP map is not fully additive as dosage-response curves show a general pattern of partial dominance (this does not contradict analytically that $m$=1, but it might reflect a possible limitation of such measure). Second, phenotype prediction using a training population is certainly population dependent. In the main text, we demonstrate that in these models, predictors arise when they are rate-limiting, that is, when they effectively limit the flux through the biomass reaction (the phenotype). Which enzyme is limiting depends on the individual and its genetic context (Fig. S9).

To find which is limiting in which individual we perform small "virtual" mutations in each enzyme sequentially (ideally they would be infinitesimal in likeliness with virtual displacements in Classical Mechanics) to identify which one incurs in growth costs (i.e., this denoting that the enzyme is limiting). In Fig. S9EF we show the results of the growth costs (colorbar) of individual virtual mutations of enzymes (rows, most rows are all 0s and irrelevant which we hid) in $10^3$ different individuals (columns) in two different populations, one in which the PGS's performance is worse than the other (Fig. S9E with $R^2$ = 0.26 and Fig. S9F with $R^2$ = 0.84). By applying an agglomerative clustering algorithm, we identify the population substructure already in $10^3$ individuals: there is a broader structure in the dendrogram of Fig. S9F than in Fig. S9E.

This explicitly demonstrates that the PGS does not only depend on the additivity of the GP map itself, but it also loses predictive power due to the integration of results of different subpopulations. The number of predictors in a population is likely a proxy of the number of subpopulations present. Therefore, the fewer the number of subpopulations the better the PGS' performance (Fig. S9C).

#### S5 Supplementary Table and Figures

| GO term | Cluster frequency | Genome frequency | p-value (corrected) | FDR | False positives |
|---|---|---|---|---|---|
| cellular BP | 100,0% | 55,6% | 5,01E-07 | 0,0% | - |
| organic substance BP | 100,0% | 58,5% | 2,76E-06 | 0,0% | - |
| BP | 100,0% | 59,2% | 4,03E-06 | 0,0% | - |
| histidine BP | 21,9% | 1,2% | 5,54E-07 | 0,0% | - |
| histidine MP | 21,9% | 1,2% | 5,54E-07 | 0,0% | - |
| lipid BP | 43,8% | 10,0% | 4,23E-05 | 0,0% | - |
| organic acid BP | 59,4% | 20,1% | 8,53E-05 | 0,0% | - |
| carboxylic acid BP | 59,4% | 20,1% | 8,53E-05 | 0,0% | - |
| small molecule BP | 68,8% | 30,0% | 5,30E-04 | 0,0% | - |
| glycerolipid BP | 21,9% | 2,5% | 5,70E-04 | 0,0% | - |
| glycerophospholipid BP | 21,9% | 2,5% | 5,70E-04 | 0,0% | - |
| lipid MP | 46,9% | 14,1% | 6,20E-04 | 0,0% | - |
| cellular lipid MP | 46,9% | 14,1% | 6,20E-04 | 0,0% | - |
| glycerophospholipid MP | 25,0% | 3,6% | 6,90E-04 | 0,0% | - |
| glycerolipid MP | 25,0% | 3,7% | 9,40E-04 | 0,0% | - |
| phospholipid BP | 25,0% | 4,0% | 1,68E-03 | 0,0% | - |
| alpha-amino acid BP | 43,8% | 13,7% | 2,56E-03 | 0,0% | - |
| phospholipid MP | 21,1% | 5,9% | 4,60E-03 | 0,0% | - |
| cellular amino acid BP | 43,8% | 14,8% | 6,37E-03 | 0,0% | - |
| GDP-mannose BP | 9,4% | 0,4% | 8,85E-03 | 0,1% | 0,02 |
| GDP-mannose MP | 9,4% | 0,4% | 8,85E-03 | 0,1% | 0,02 |
| long-chain fatty acid BP | 9,4% | 0,4% | 8,85E-03 | 0,1% | 0,02 |

**Table S1. GO enrichment analysis of predictors with the largest effect sizes.** We further confirm that the top genetic predictors cluster into only a few biosynthetic and metabolic processes (BP and MP, respectively). They are mainly related with amino acids, phospholipids, fatty acids and mannose (Cherry et al., 2012).

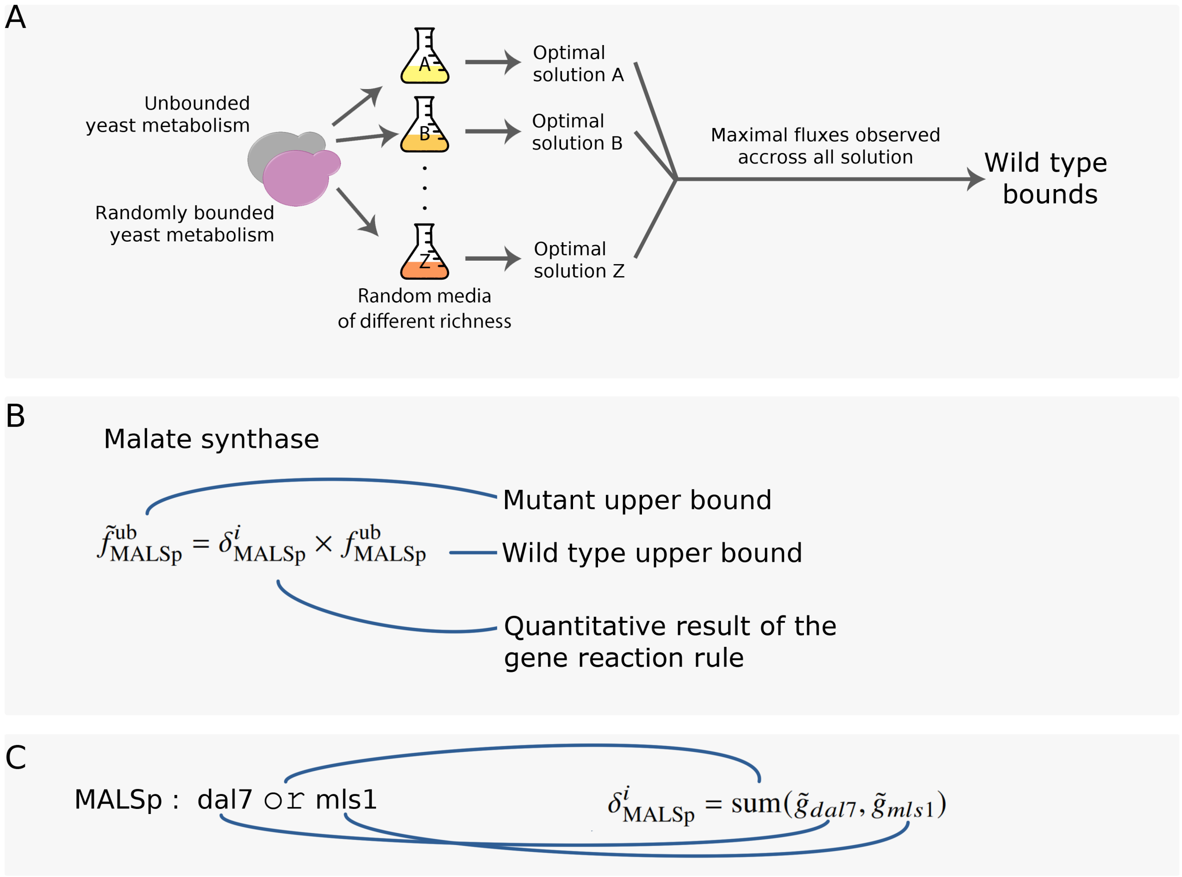


**Figure S1. Modeling of quantitative mutations.** We characterize mutations by a decrease in enzyme efficiency with respect to a wild type, "reference", or "maximum" value. (**A**) To find the wild type bounds, we expose the yeast metabolism to a series of environmental and genetic adaptations and compute the maximum fluxes observed in the solutions. Specifically, we compute pairs of optimal solutions in $10^4$ random media from a totally unbounded and a randomly bounded yeast metabolisms (random bounds change in every medium; Methods). (**B**) Then, the bounds for a given reaction of a given mutant is the product of the wild type bounds and a fractional value resulting from the quantitative interpretation of gene reaction rules. (**C**) These rules describe how enzymes control the reaction. Namely, isozymes (and coenzymes) are modeled by an "OR" (and an "AND") operator which we translate by the sum (and the minimum) of gene dosages. The "AND" means that both enzymes are necessary, and thus the minimum dosage of both will be the limiting factor; in contrast, the "OR" operator means that either enzyme can carry out the reaction, hence we use the sum of gene dosages. We repeat this process iteratively until the entire reaction rule is translated into a fractional bound.

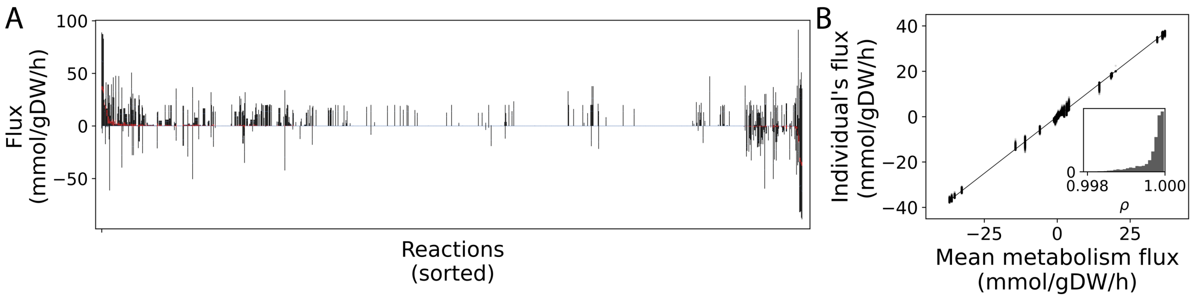


**Figure S2. Genetic variation leads to flux variability, which is well described by the mean metabolism.** (**A**) Fluxes that are accessible in the population, i.e. maximal bounds (vertical black lines) and range of values observed in the population (vertical red lines) for each reaction (x-axis). Blue lines represent 70% of such maximal bounds, which is approximately the largest restriction in the default population (with dosages sampled from a normal distribution with unit mean and $\sigma = 0.1$). We find that the genetic variation with which the population was generated leads to variability in some solution fluxes of the individuals, which ultimately translate into growth variability. (**B**) Despite this variability in solution fluxes, we can define a "mean" metabolism in which the flux through each reaction is the observed mean across the population. Black dots depict data of the reactions of all individuals in the population, and the inset shows the distribution of linear correlations between each individual's solution and the mean metabolism.

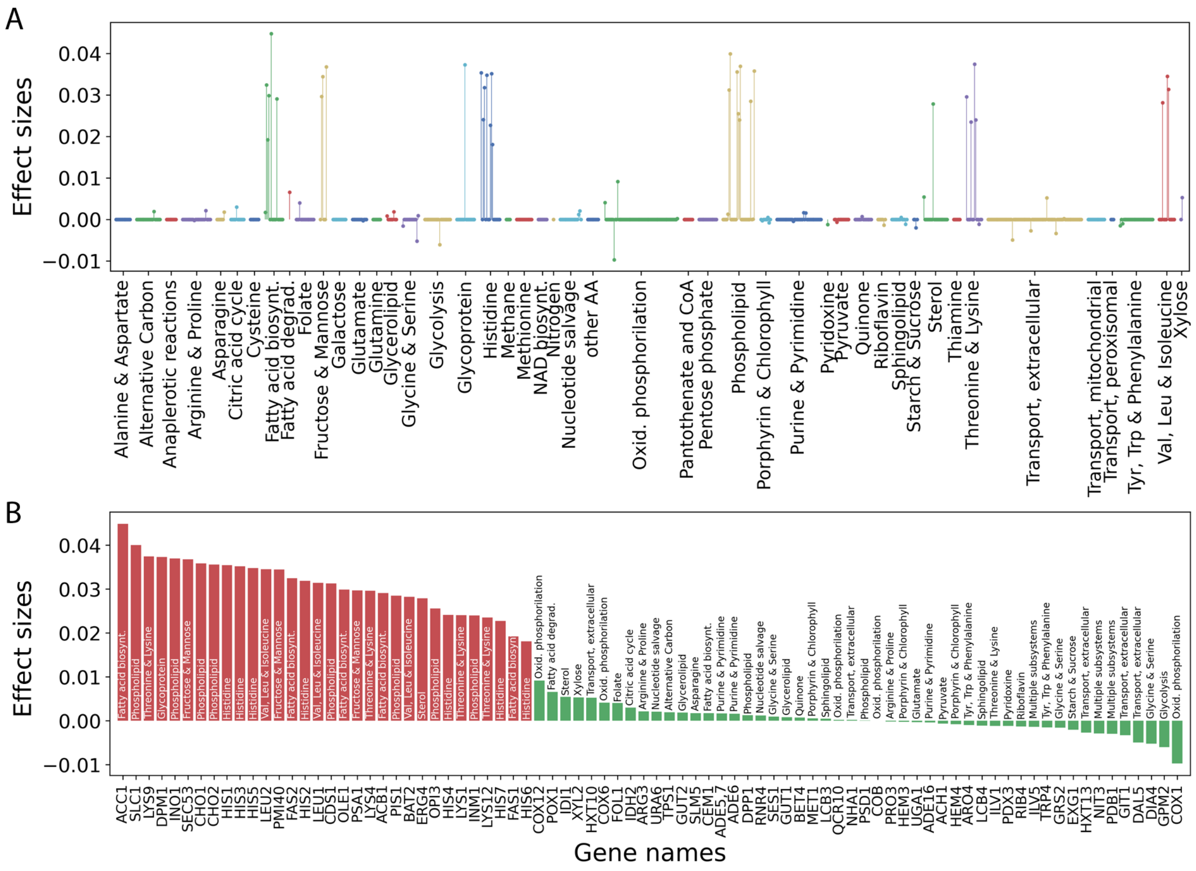

**Figure S3. Genetic predictors belong to a handful of metabolic subsystems.** (**A**) Manhattan-like plot showing the effect sizes (y-axis) of genes grouped by yeast metabolic subsystems (x-axis; arbitrary colors). We find that genes with large effect sizes belong to a handful of subsystems related to protein synthesis, cell membrane and organelle compartmentalization. (**B**) Effect sizes of all predictors identified in the PGS of the Results section 2 (colors as in Fig.2) and their corresponding metabolic subsystem.

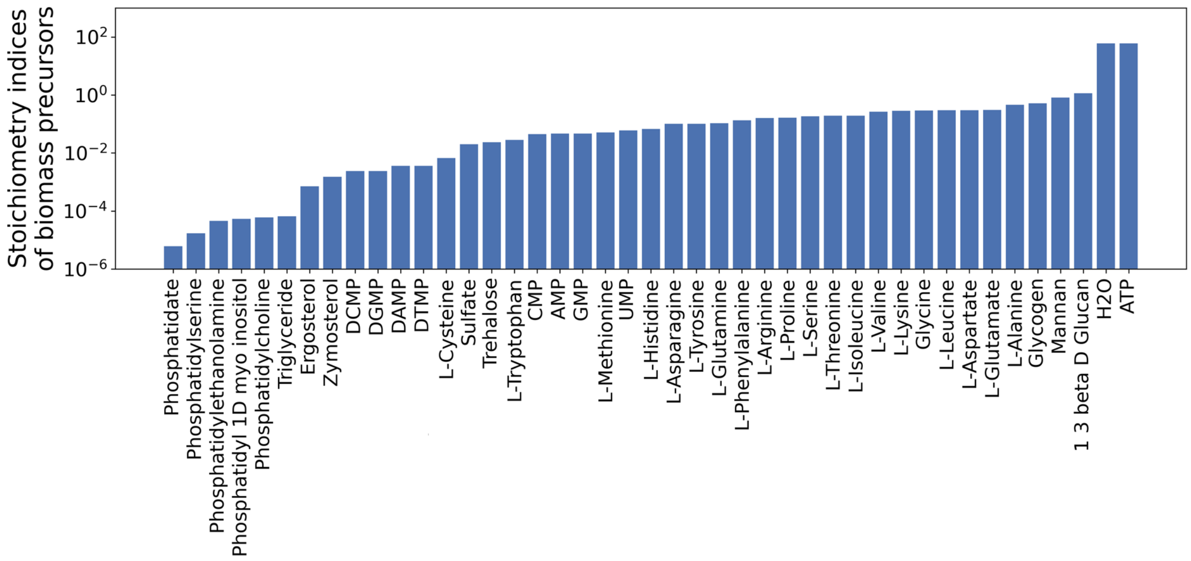


**Figure S4. Biomass precursors and their stoichiometric coefficients in the biomass reaction.** The biomass reaction involves 43 precursor metabolites (x-axis) but with stoichiometric coefficients spanning several orders of magnitude (y-axis, in log scale). For example, the most consumed precursors are ATP and water.

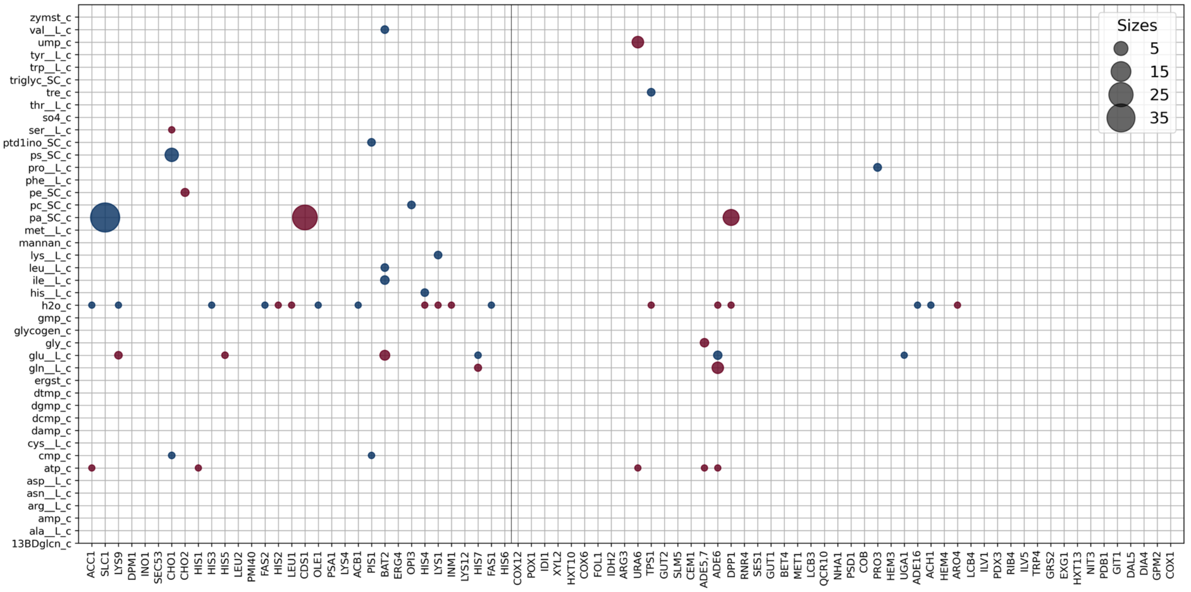


**Figure S5. Contribution of genetic predictors to the mean production or depletion of biomass precursors.** Each genetic predictor (x, axis; sorted by effect size) participates in a number of reactions that might involve biomass precursors (y-axis). We here show the mean consumption (red circles) or production (blue circles) across the entire population ($10^4$ individuals). Circle sizes are proportional to the absolute value of the mean contribution relative to the biomass consumption. Figure at full size available online (**?**).

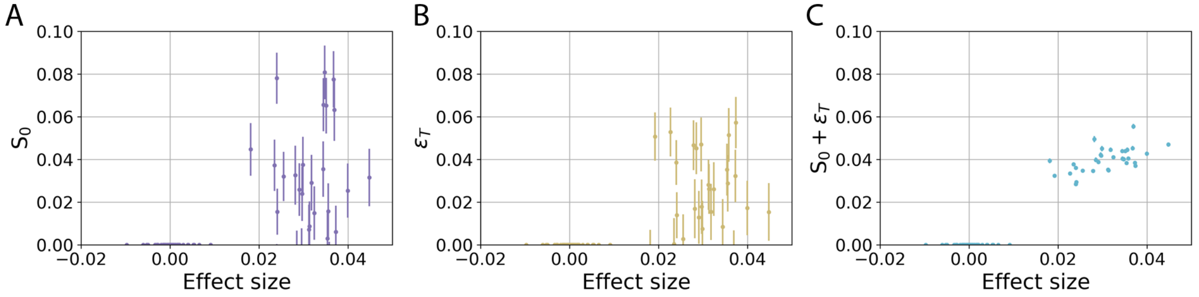


**Figure S6. Effect sizes correlate with global sensitivity indices.** We show (**A**) the first order index $S_0$, (**B**) the total epistasis $\epsilon_T$ and (**C**) the total effects $S_0 + \epsilon_T$ as a function of effect size (Methods). The linear correlations among all genes are $\rho^{all}_{S_0} = 0.63$, $\rho^{all}_{\epsilon_T} = 0.48$ and $\rho^{all}_{+} = 0.98$ respectively; or among only large effect predictors $\rho^{pred}_{S_0} = 0.19$, $\rho^{pred}_{\epsilon_T} = -0.08$ and $\rho^{pred}_{+} = 0.57$. We show the mean values and a standard deviation of $> 10^6$ simulations for each gene (Methods).

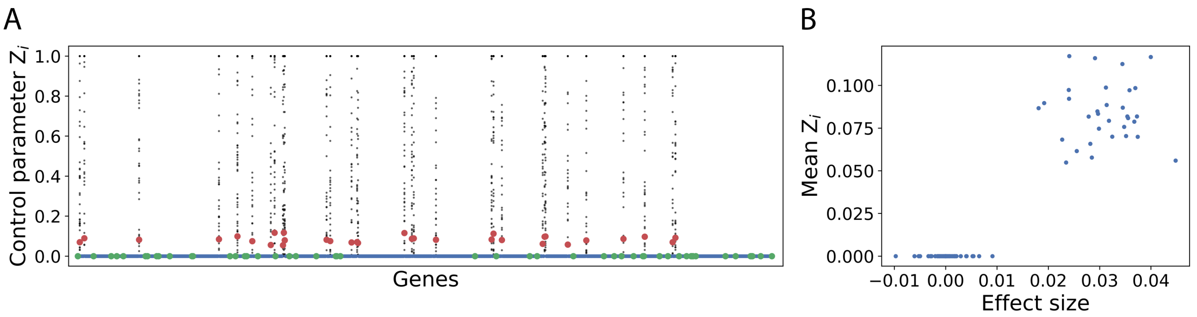


**Figure S7. Control parameters $Z_i$ of growth.** (**A**) Control parameters quantify the relative impact of individual changes of a gene's dosage on growth, and they are highly context-dependent. We show individual values, for every genetic background (black dots), and the mean values across the population (colored dots, colors as in Figure 2). (**B**) Mean values of the control parameters across different backgrounds anticipates predictor genes within a PGS.

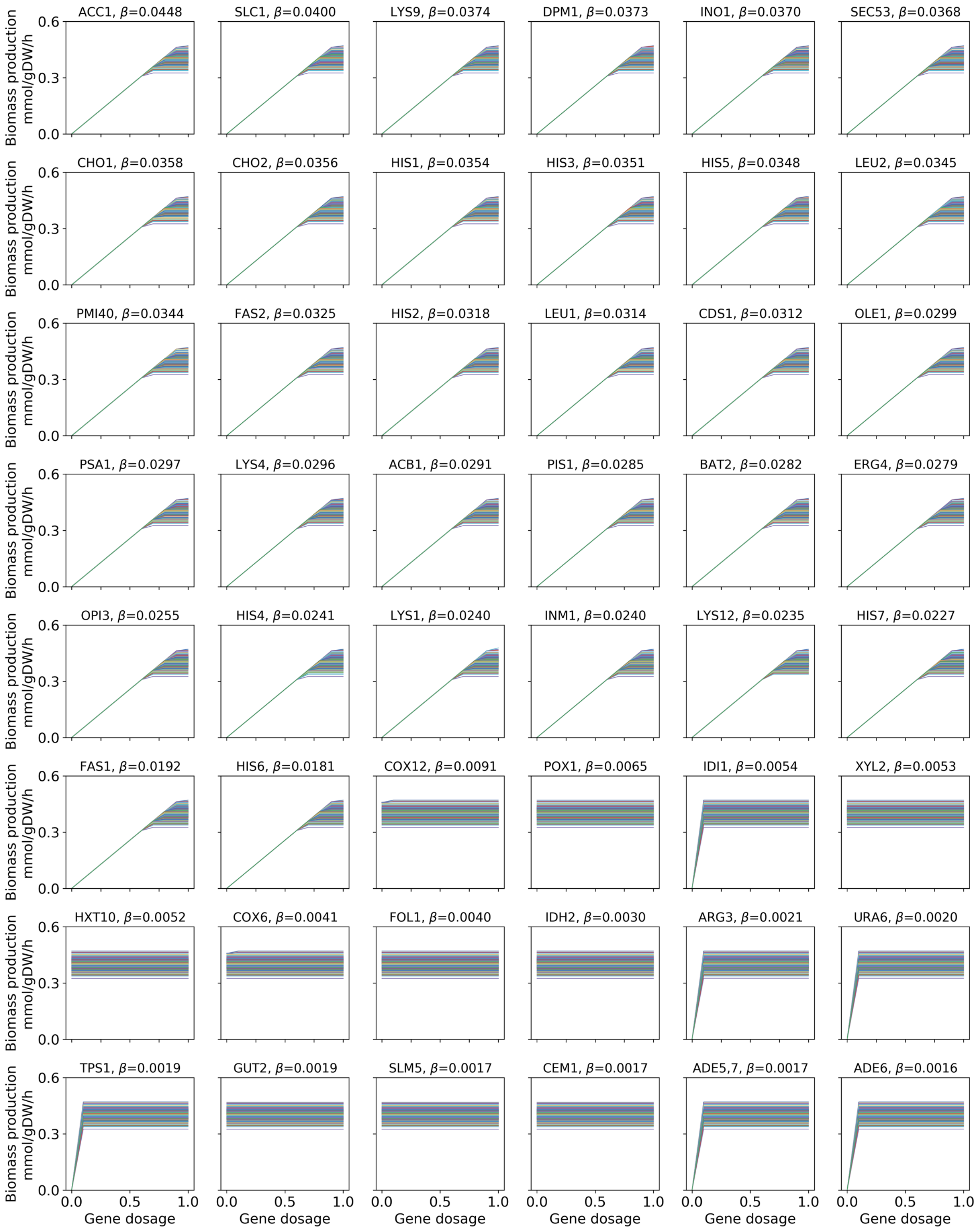


**Figure S8. Dosage-response profiles of all predictor genes.** We computed the dosage-response profiles for all predictor genes in 200 genetic backgrounds by individually tuning the corresponding dosage from $g = 0$ to $g = 1$ and computing the growth rate with FBA. Observe that i) all top predictors (with $\beta > 0.01$) are essential. That is, growth is null if $g = 0$; ii) that only top predictors display a recurrent dosage-response profile and that iii) the profiles of genes with $\beta < 0.01$ are constant in the range mostly accessed by the population $0.7 < g < 1$.

[H]

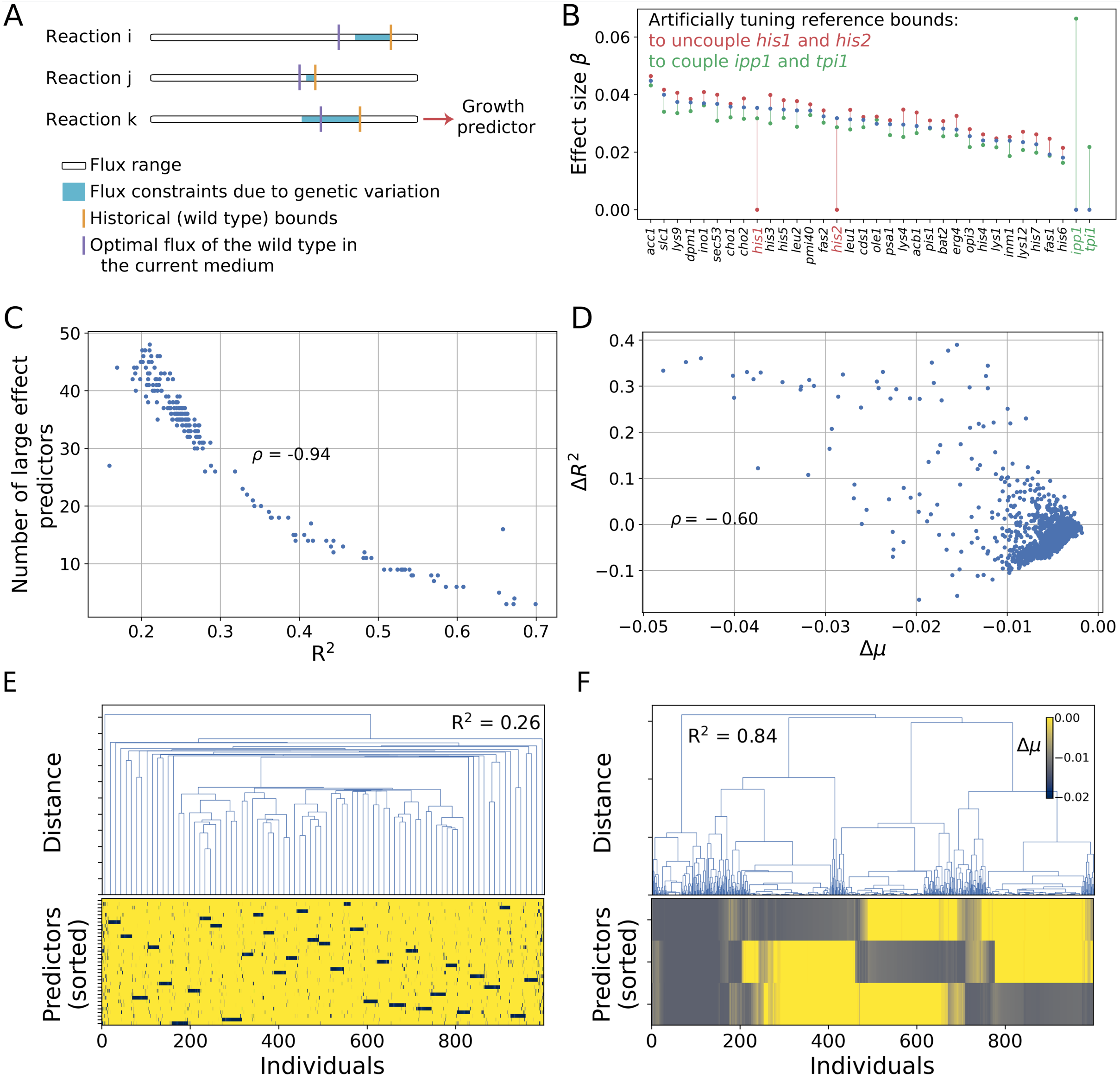


*(Caption on next page.)*

### References


Cherry, J. M., Hong, E. L., Amundsen, C., Balakrishnan, R., Binkley, G., Chan, E. T., Christie, K. R., Costanzo, M. C., Dwight, S. S., Engel, S. R., Fisk, D. G., Hirschman, J. E., Hitz, B. C., Karra, K., Krieger, C. J., Miyasato,

**Figure S9. Genetic variation and the optimal functional mode in the growth medium determine the predictor character of a gene in metabolism.** (**A**) Cartoon of three different reactions depicting the flux range available in the wild type (white), and their optimal flux in the current growth medium (purple; functional mode). In a population, genetic variation translates into a range of flux constraints (cyan) but it can become cryptic if it does not constrain the optimal flux or if gene reaction rules filter it out as in reactions $i$ and $j$. Only if genetic variation limits the flux of the functional mode, its associated gene(s) can become growth predictors (as in reaction $k$). (**B**) We confirmed this in two experiments. In the first, we artificially increase the wild type bounds of reactions associated to *his1* and *his2*. In the second, on the contrary, we decrease the bounds of reactions controlled by genes *ipp1* and *tpi1*. In the former and latter experiments, we successfully uncoupled and coupled the genes to growth prediction, respectively. We show the effect sizes of large effect predictors in the original PGS (blue) and after both experiments (red and green, respectively). (**C**) If we iterate the previous experiment by randomly selecting between 10 and 60 reactions, and randomly (un)coupling them from growth prediction, we find that the performance of a PGS correlates negatively with the number of large effect predictors (linear $\rho = -0.94$). Thus, the more growth-limiting reactions, the more predictors and the worse is predictability within a PGS. (**D**) In another experiment, we study the impact of "disabling" the metabolism to activate new reactions that are inactive in the wild type solution. This leads to a better performance of a PGS ($\Delta R^2$) at the expense of fitness costs ($\Delta\mu$), which negatively correlate (linear $\rho = -0.60$). (**E-F**) Growth costs (colorbar) of individual virtual mutations of enzymes identify the structure of limiting reactions in two populations for which the PGS performance differs ($R^2 = 0.26$ and $R^2 = 0.84$ in E and F respectively). Enzyme mutations in rows, $10^3$ different individuals in columns. There is a simpler structure in the dendrogram of panel F (with larger $\mathrm{R}^2$) than in panel E (with smaller $\mathrm{R}^2$).


S. R., Nash, R. S., Park, J., Skrzypek, M. S., Simison, M., Weng, S., and Wong, E. D. (2012). "Saccharomyces genome database: the genomics resource of budding yeast." *Nucleic Acids Res.*, 40(Database issue), D700–5.

Gjuvsland, A. B., Vik, J. O., Woolliams, J. A., and Omholt, S. W. (2011). "Order-preserving principles underlying genotype–phenotype maps ensure high additive proportions of genetic variance." *Journal of Evolutionary Biology*, 24(10), 2269–2279.

Gjuvsland, A. B., Wang, Y., Plahte, E., and Omholt, S. W. (2013). "Monotonicity is a key feature of genotype-phenotype maps." *Frontiers in Genetics*, 4.

Hill, W. G., Goddard, M. E., and Visscher, P. M. (2008). "Data and theory point to mainly additive genetic variance for complex traits." *PLOS Genetics*, 4(2), 1–10.

Kacser, H. and Burns, J. A. (1981). "The molecular basis of dominance." *Genetics*, 97(3-4), 639–666.

Kacser, H., Burns, J. A., Kacser, H., and Fell, D. A. (1995). "The control of flux." *Biochemical Society Transactions*, 23(2), 341–366.

Nobile, M. S., Coelho, V., Pescini, D., and Damiani, C. (2021). "Accelerated global sensitivity analysis of genome-wide constraint-based metabolic models." *BMC bioinformatics*, 22(Suppl 2), 78–78.

Orth, J. D., Thiele, I., and Palsson, B. (2010). “What is flux balance analysis?.” *Nature Biotechnology*, 28(3), 245–248.

Saltelli, A., Annoni, P., Azzini, I., Campolongo, F., Ratto, M., and Tarantola, S. (2010). “Variance based sensitivity analysis of model output. design and estimator for the total sensitivity index.” *Computer Physics Communications*, 181(2), 259–270.

Saltelli, A., Ratto, M., Andres, T., Campolongo, F., Cariboni, J., Gatelli, D., Saisana, M., and Tarantola, S. (2008). *Global Sensitivity Analysis: The Primer*. Wiley.

Sobol, I. M. (1993). “Sensitivity analysis for non-linear mathematical models.” *Mathematical modelling and computational experiment*, 1, 407–414.

Sobol, I. M. (2007). “Global sensitivity indices for the investigation of nonlinear mathematical models.” *Matematicheskoe Modelirovanie*, 19(11), 23–24.
